# Engineering Brain Parasites for Intracellular Delivery of Therapeutic Proteins

**DOI:** 10.1101/481192

**Authors:** Shahar Bracha, Karoliina Hassi, Paul D. Ross, Stuart Cobb, Lilach Sheiner, Oded Rechavi

**Author notes:** Equal contribution (Lilach Sheiner, Oded Rechavi).

## Abstract

Protein therapy has the potential to alleviate many neurological diseases; however, delivery mechanisms for the central nervous system (CNS) are limited, and intracellular delivery poses additional hurdles. To address these challenges, we harnessed the protist parasite *Toxoplasma gondii,* which can migrate into the CNS and secrete proteins into cells. Using a fusion protein approach, we engineered *T. gondii* to secrete therapeutic proteins for human neurological disorders. We tested two secretion systems, generated fusion proteins that localized to *T. gondii*’s secretory organelles and assessed their intracellular targeting in various mammalian cells including neurons. We show that *T. gondii* expressing GRA16 fused to the Rett syndrome protein MeCP2 deliver a fusion protein that mimics the endogenous MeCP2, binding heterochromatic DNA in neurons. This demonstrates the potential of *T. gondii* as a therapeutic protein vector, which could provide either transient or chronic, *in situ* synthesis and delivery of intracellular proteins to the CNS.

## Introduction

Protein therapeutics have the potential to treat various human disorders: many genetic diseases caused by mutations in a specific protein can be rescued by delivery of the functional protein, treating the root of the disease. In addition, proteins with downstream or compensatory activity can provide indirect rescue or mitigation of disease symptoms (Leader, Baca, & Golan, 2008). However, owing to their macromolecular nature, the delivery of therapeutic protein to target tissues is extremely challenging. Low functional stability and rapid loss of activity following administration or during storage, along with low permeability through physiological barriers, limit the delivery of active proteins (Leader et al., 2008). Efficient delivery is particularly challenging for neurological diseases due to the blood-brain barrier (BBB), which tightly regulates the transport of molecules to the brain, and blocks the transport of most large, charged and hydrophilic molecules (DiNunzio & Williams, 2008). In addition to specific delivery to the brain, intracellular delivery of proteins poses additional hurdles. Passive delivery is often precluded as most proteins are unable to spontaneously penetrate through the cell membrane or become endocytosed effectively (Torchilin, 2008). Furthermore, intracellular delivery has been proven especially difficult for neuronal cells (Karra & Dahm, 2010).

The use of parasites that co-exist with humans for the treatment of chronic and degenerative diseases has a long history. However, these treatments mostly rely on harnessing the ability of the parasite to suppress or induce specific immune responses; examples include the use of immunosuppressive helminths (parasitic worms) for treating allergies and autoimmune diseases (Helmby, 2015); the pre-antibiotics fever therapy for neurosyphilis (Whitrow, 1990) and the use of Bacillus Calmette-Guerin bacteria for cross-vaccination and more recently cancer immunotherapy (Fuge, Vasdev, Allchorne, & Green, 2015). In addition, the rise of synthetic biology offers new methods for bioengineering organisms that mediate designed therapeutic intervention. Some notable examples include metabolically engineered microbiome bacteria (Lu, Mimee, Citorik, & Pepper, 2017); antigen-presenting microorganisms that induce immune responses against targeted pathogens or tumor cells (Wood & Paterson, 2014); and the use of engineered viruses for gene therapy (Colella, Ronzitti, & Mingozzi, 2018).

*Toxoplasma gondii* is a highly prevalent eukaryotic parasite of the phylum Apicomplexa. Through co-evolution with its hosts, *T. gondii* acquired sophisticated mechanisms to migrate into the CNS and establish quiescent cysts that can persist for the lifetime of the host (Hill, Chirukandoth, & Dubey, 2005; Mendez & Koshy, 2017). The primary hosts of *T. gondii* are felids, but it can infect all warm-blooded organisms as intermediate hosts. In immunocompetent humans, infections are typically asymptomatic or with only mild, short lived flu-like symptoms. As a result, most infections go unnoticed, and it is estimated that about a third of the human population is chronically infected with *T. gondii* (Pittman & Knoll, 2015)*. T. gondii*’s most common route of entry is by ingestion of infective cysts. Upon entry into the human body, it actively migrates to the brain and passes the BBB by three putative mechanisms: infecting and hitch-hiking on immune cells that infiltrate the CNS through a “trojan horse” mechanism, migrating through the tight junctions of the BBB, or invading into the endothelial cells of the BBB and egressing into the basolateral side (Mendez & Koshy, 2017). In the brain, *T. gondii* interacts and resides mostly in neurons (Cabral et al., 2016). Importantly, to survive inside the host, *T. gondii* secretes a myriad of effector proteins into the host cells and into the extracellular environment (Hakimi, Olias, & Sibley, 2017).

These characteristics led us to propose the use of *T. gondii* as a biological vector for the delivery of therapeutic proteins to the CNS. *T. gondii*’s active migration can provide high specificity to the CNS, and its specialized secretion systems can be used for extracellular or intracellular protein delivery. Here we describe the development of this system by engineering transgenic *T. gondii* lines that synthesize and deliver therapeutic proteins into mammalian cells. Our work demonstrates that engineered *T. gondii* is a promising biological vector for mediating protein therapy for neurological diseases.

## Results

### General approach for engineering *T. gondii* to secrete heterologous proteins

Our approach for engineering *T. gondii* to secrete heterologous proteins was based on fusing therapeutic proteins to endogenously-secreted proteins of the parasite (Fig. 1). We exploited two *T. gondii* secretion systems that secrete proteins into cells: rhoptries, which inject proteins directly into the cytosol of cells before and during cell invasion (Boothroyd & Dubremetz, 2008); and dense granules, which secrete proteins after the parasite invaded and resides inside the intracellular parasitophorous vacuole (PV). Dense granule proteins can be secreted continuously throughout *T. gondii*’s persistence inside the cell. However, in order for such proteins to reach the host cell from within the PV, they must subsequently be exported through the PV membrane (PVM) and released into the host cell cytosol (Hakimi et al., 2017).

**Fig. 1.**
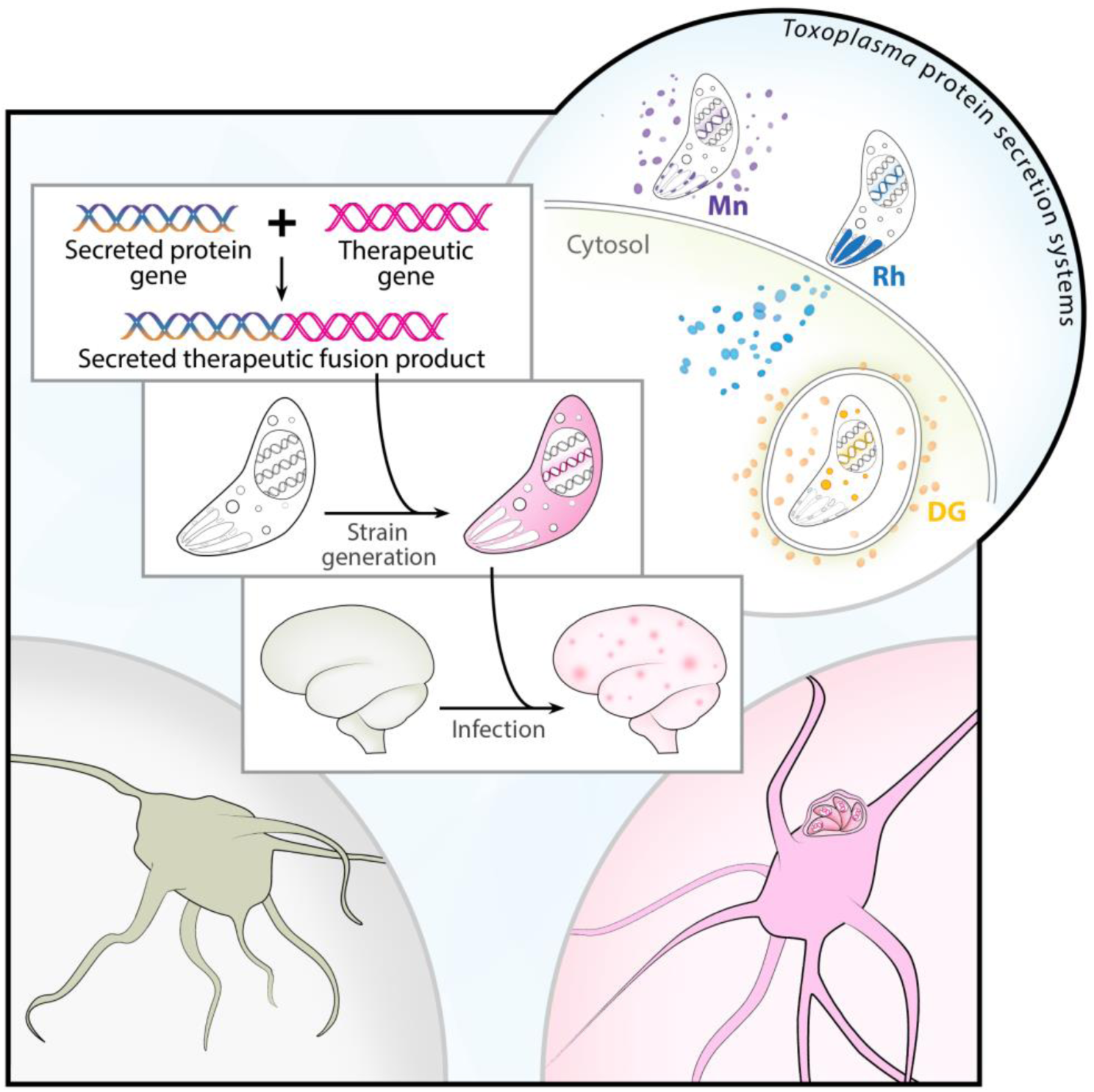
Generating *T. gondii* for delivery of heterologous therapeutic proteins. A therapeutic protein of interest is genetically fused to an endogenously-secreted protein of the parasite, which targets it for secretion from the respective secretory organelle. The genetic construct encoding for the fusion protein is introduced into *T. gondii* to generate transgenic lines which secrete the therapeutic protein. The transgenic *T. gondii* actively migrates to the CNS, bypasses the BBB and delivers the protein to neurons. The top right corner presents the three secretion systems of *T. gondii*, which secrete proteins using different mechanisms and at different stages of cell invasion or intracellular persistence. Mn-microneme, Rh-rhoptry, DG-dense granule.

To test these two secretion systems, we selected a series of human and mouse proteins which (a) have validated therapeutic effects in models of human neurological diseases, (b) are required in neurons, (c) are required in relatively low levels, and preferentially (d) have robust *in vitro* activity assays compatible with *T. gondii* culturing. We also aimed to select proteins of various sizes, target cellular localizations and cellular functions; this allowed us to assess the effect of these protein features on targeting in the parasite, ability to be secreted by the rhoptry or dense granules, and export from the PV. According to these criteria, we selected the following proteins: Aspartoacylase (*ASPA*, 36kDa), a nuclear-cytosolic enzyme associated with Canavan disease (Leone et al., 2012); Survival of Motor Neuron (*SMN1*, 32kDa), a nuclear-cytosolic protein associated with spinal muscular atrophy (Bowerman et al., 2017); Glial Derived Neurotrophic factor (*GDNF*, 24kDa), a signaling peptide associated with neuroregeneration and a variety of neurodegenerative diseases (Allen, Watson, Shoemark, Barua, & Patel, 2013); Parkin (*PARK2*, 42kDa), a nuclear-cytosolic E3 ubiquitin-ligase associated with Parkinson’s disease (Mochizuki, 2007); Galactosylceramidase (*GALC*, 77kDa), a lysosomal enzyme associated with Krabbe disease (Rafi, Rao, Luzi, Curtis, & Wenger, 2012), Transcription factor EB (*TFEB*, 53kDa), a nuclear-cytosolic transcription factor associated with a variety of neurodegenerative and lysosomal storage diseases (Napolitano & Ballabio, 2016); and Methyl-CpG binding Protein 2 (*MECP2*, 54kDa), a nuclear DNA-binding protein associated with Rett syndrome (Katz et al., 2016). To examine the effect of codon usage on expression levels and targeting of the heterologous proteins, the mammalian genes *ASPA*, *GALC* and *MECP2* were tested in both their native mammalian sequence and codon optimized according to *T. gondii*’s codon usage (labeled ‘opt’, e.g. *MECP2opt*). The lysosomal *GALC* was tested in an additional variation containing a TAT protein transduction domain, which promotes protein transduction across membranes and improves cross-correction between cells (Meng, Eto, Schiffmann, & Shen, 2013).

### Targeting proteins to the rhoptry secretory organelle for transient delivery

The first secretion system we tested was the rhoptries. As some rhoptry proteins are injected before cell invasion, their secretion does not depend on productive invasion and persistence of the parasite inside the host cell. Furthermore, quantification in the CNS of mice showed that rhoptry proteins can be injected by *T. gondii* into 30-50 times more cells in the brain than those productively infected (Koshy et al., 2012). This suggests an advantage of rhoptry secretion for transient delivery of proteins, as each parasite can secrete rhoptry proteins into multiple cells without invading them.

In order to co-opt rhoptry secretion, heterologous proteins were fused to the rhoptry protein Toxofilin, which is injected into the cytosol of host cells upon initial parasite engagement (Boothroyd & Dubremetz, 2008). The fusion to Toxofillin was modelled after an ingenious reporter line of *T. gondii* expressing Toxofilin-fused Cre recombinase, which enabled the labeling of cells contacted by the parasite both *in vitro* and *in vivo* (Cabral et al., 2016; Koshy et al., 2010; Lodoen, Gerke, & Boothroyd, 2010). We generated parasites with genomic integration of constructs encoding for Toxofilin fused to different variations of: ASPA, SMN, GDNF, PARK2, GALC, MECP2, and TFEB (**Table S1**). Expression was controlled by the endogenous promoter of *Toxofilin*, and an HA tag was added for immunolabeling of the fusion proteins. Most fusion proteins mis-localized to other organelles of the parasite, predominantly to the endoplasmic reticulum (ER), golgi and micronemes. However, Toxofilin-fused GDNF, PARK2 and codon-optimized TFEB (*TFEBopt*) were successfully localized to the rhoptries (Fig. 2, **Table S1**).

**Fig. 2.**
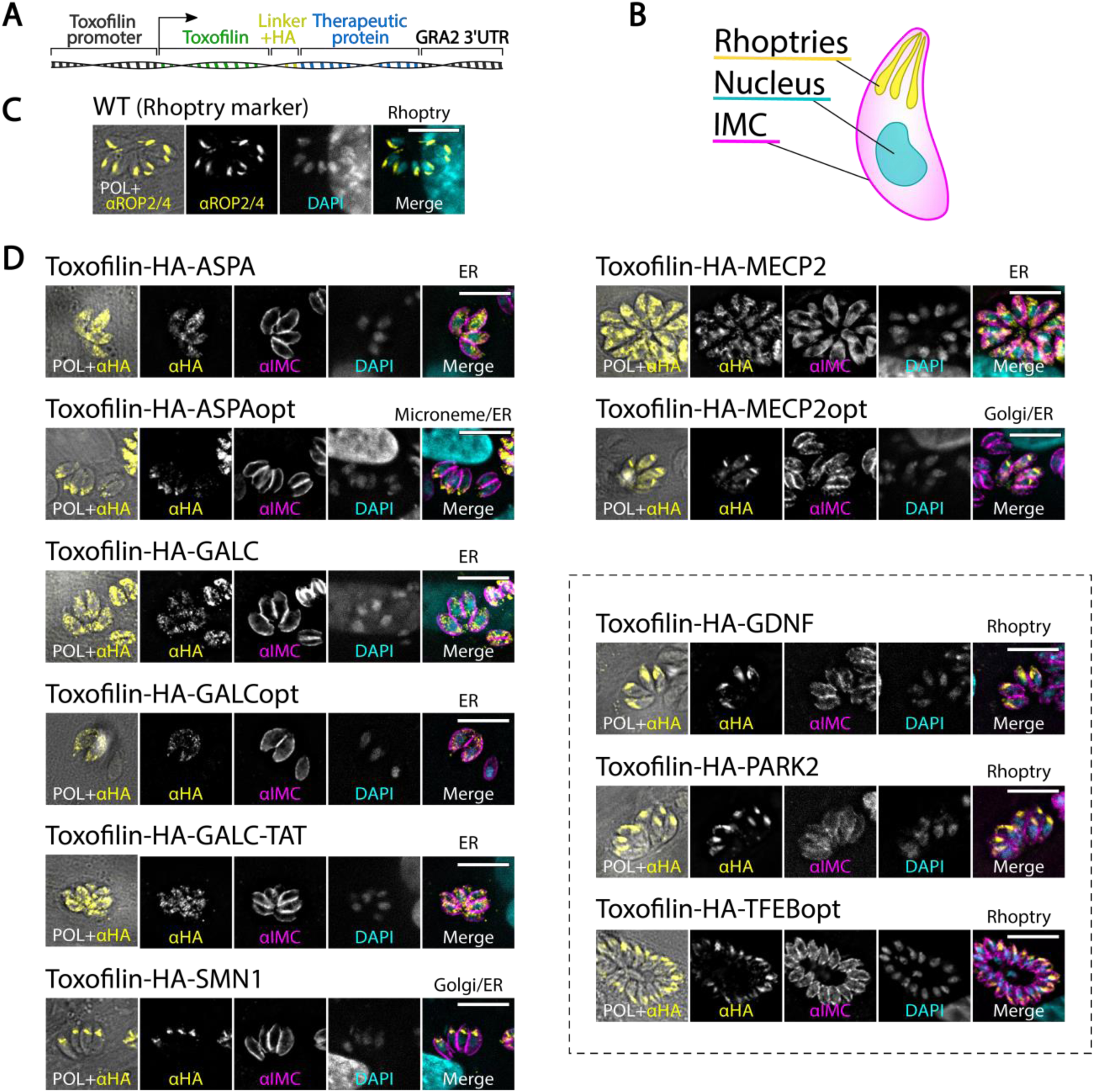
Targeting of therapeutic proteins to *T. gondii*’s rhoptries by fusion to Toxofilin. (**A**) Scheme of the genetic construct used to generate *T. gondii* expressing Toxofilin-fused therapeutic proteins. (**B**) Illustration of a *T. gondii* parasite. IMC-inner membrane complex, marking the parasite outline. (**C**) Intracellular *T. gondii* immunostained with the rhoptry marker anti-ROP2/4. (**D**) Intracellular *T. gondii* stably expressing different Toxofilin-fused proteins related to human neurological diseases. Selected images represent localization of the proteins to the rhoptries (TFEBopt, PARK2, GDNF) or in the lack of any, the most common localizations in the stable pool (full list of observed localizations in Table S1). Text above the merge images indicates the localization of the fusion protein in the image. Scale bar: 10µm.

### Targeting proteins to the dense granule secretory organelle for continuous delivery

As rhoptry secretion occurs as a discrete event upon initial parasite engagement, it provides only transient protein delivery. In contrast, dense granule proteins can be secreted continuously as the parasite persists inside a cell. Therefore, we assessed dense granule-mediated secretion for continuous delivery of intracellular proteins. For this purpose, we designed translational fusions to GRA16. GRA16 is secreted from the dense granules to the parasitophorous vacuoles (PV), exported from the PV to the host cell cytosol, and localizes to the host cell nucleus (Bougdour et al., 2013). We generated parasites with genomic integration of constructs encoding for GRA16 fused to different variants of: ASPA, SMN, GALC, MECP2, and TFEB (**Table S2**). Expression was controlled by the endogenous promoter of *GRA16*, and an HA tag was added for immunolabeling of the fusion proteins.

GRA16-fused GALC-TAT was not expressed in *T. gondii*. GRA16-fused GALCopt was expressed, but localized only to the ER of the parasites and was not secreted to the PV, nor detected in the host cell. ASPA, ASPAopt and GALC fused to GRA16 localized to the PV but were not detected in the host cell, indicating that they were successfully targeted to the dense granules and secreted to the PV, but not exported in detectable levels. Importantly, the GRA16-fused nuclear proteins MeCP2 (*MECP2opt*, total fusion protein size 110kDa), SMN (*SMN1*, 88kDa) and TFEB (*TFEBopt*, 109kDa) localized to both the PV and the host cell nucleus. This indicates that these proteins were successfully targeted to the dense granules, secreted to the PV, exported to the host cell cytosol and accumulated in their site of activity, the host nucleus (**Fig. 3**) (Bowerman et al., 2017; Katz et al., 2016; Napolitano & Ballabio, 2016).

Interestingly, by analyzing the level of intrinsic disorder for each of the tested fusion proteins, we found a strong correlation (R=0.975, P=4*10-5) between the level of localization to the host cell nucleus and the average intrinsic disorder score of the fused protein (based on IUPred2A (Mészáros, Erdős, & Dosztányi, 2018)) (Fig. 3F, Fig. S1). This provides strong support for involvement of protein structural disorder in export through the PVM, and for the hypothesized translocon model of protein export (Hakimi & Bougdour, 2015; Hakimi et al., 2017).

**Fig. 3.**
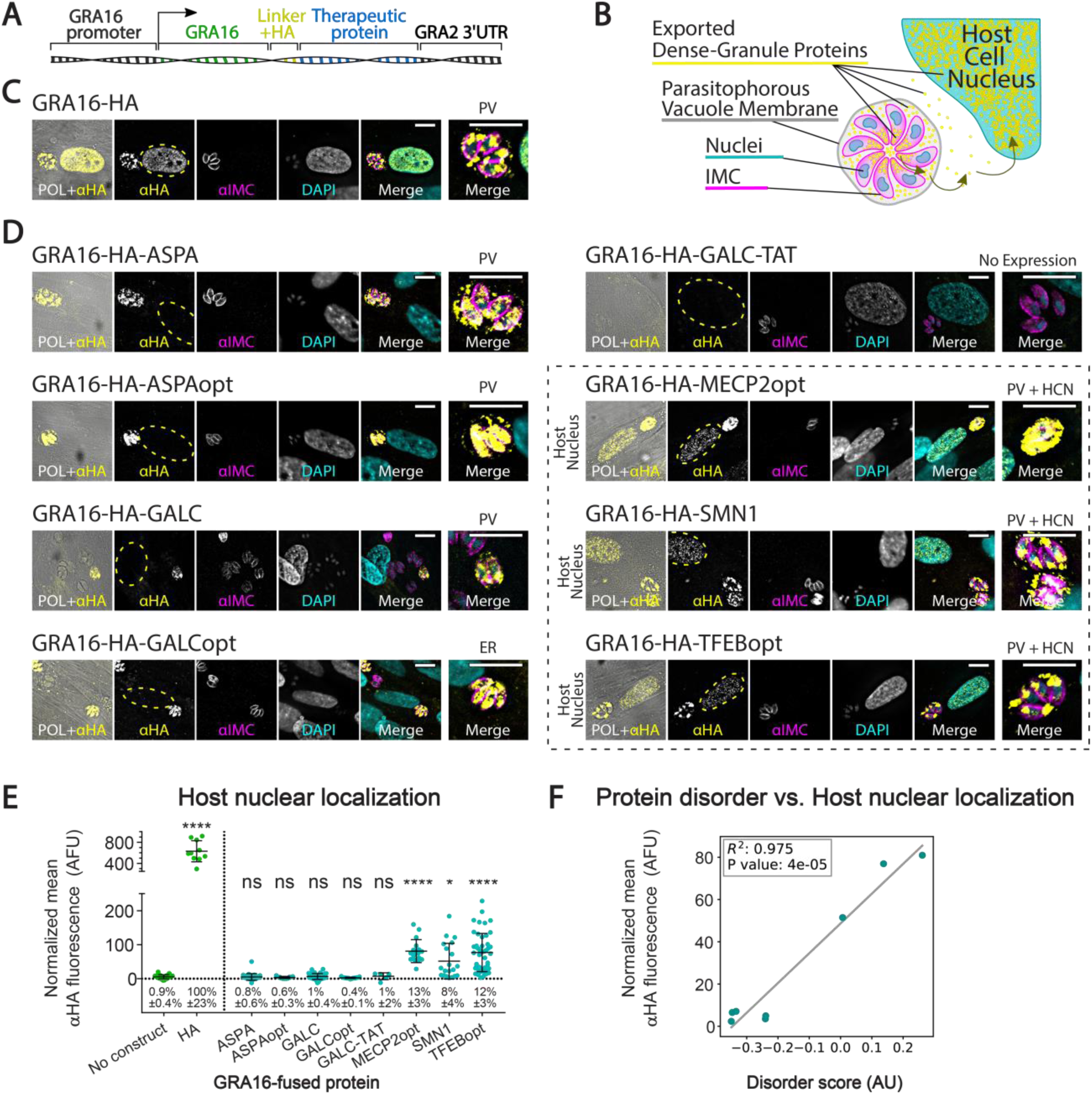
Targeting of therapeutic proteins to *T. gondii*’s dense granules, parasitophorous vacuole and host cell nucleus by fusion to GRA16. (**A**) Scheme of the genetic construct used to generate *T. gondii* expressing GRA16-fused therapeutic proteins. (**B**) Illustration of an intracellular parasitophorous vacuole containing eight *T. gondii* parasites. Yellow-exported dense-granule protein targeted to the nucleus of the host cell. (**C-D**) Intracellular *T. gondii* stably expressing HA-tagged GRA16 (**C**) or different GRA16-fused proteins related to human neurological diseases (**D**). Selected images represent localization of the proteins to the PV and host cell nucleus (TFEBopt, MECP2opt, SMN1), to the PV alone (ASPA, ASPAopt, GALC) or in the lack of any, the most common localizations in the stable pool (full list of observed localizations in Table S2). Rightmost image of each set shows a close-up of the parasite vacuole, and the text above it indicates the localizations of the fusion protein in the image. Yellow dashed lines mark the host cell nucleus (HCN). (**E**) Fluorescence quantification of the anti-HA signal in the nucleus of infected host cells. Error bars represent mean ±SD. N=6-47 cells per condition. Take note that the y-axis is divided to two segments, to account for the high values recorded from GRA16-HA. AFU-arbitrary fluorescence units. Text below each scatter plot shows the mean relative nuclear localization ±95% confidence interval (relative to GRA16-HA). Significance represents the difference between the fusion protein and the parental strain (‘no construct’), calculated by one-way ANOVA with multiple comparisons (Dunnett test). **** P<0.0001; * P<0.0332; ns-not significant. (**F**) Correlation between host nuclear localization and the level of intrinsic disorder of the fused therapeutic protein, based on IUPred2A (see Fig. S1). Linear regression line drawn in grey. Scale bar: 10µm.

To assess whether a shorter fragment of GRA16 could replace the full-length protein as a carrier for secretion, we tested three truncated variants of GRA16 fused to MeCP2. However, as none of the truncated fusion proteins were detected in the host cell nucleus, we decided to continue with the full-length GRA16 as the carrier protein (Fig. S2).

Since GRA16-fused MeCP2 and TFEB displayed the most robust delivery and targeting to the host cell nucleus, we focused on these fusion proteins for the rest of the study. The *T. gondii* lines expressing these proteins were named GRA16-MeCP2 (expressing *GRA16-HA-MECP2opt*) and GRA16-TFEB (expressing *GRA16-HA-TFEBopt*). In addition, *T. gondii* expressing HA-tagged GRA16 which is not fused to any protein (GRA16-HA) was generated as a control.

### Kinetics of GRA16-mediated protein delivery in human fibroblasts and neurons

To characterize the kinetics of GRA16-mediated protein delivery *in vitro*, we used high-content imaging to quantify infection rate and nuclear protein delivery for parasites expressing GRA16 alone and GRA16 fused to MeCP2 or TFEB. We infected human foreskin fibroblast (HFF) cells using different MOI (Multiplicity Of Infection-the proportion of parasite per host cell administered to the culture) and followed nuclear fluorescence over time (Fig. 4A-C).

**Fig. 4.**
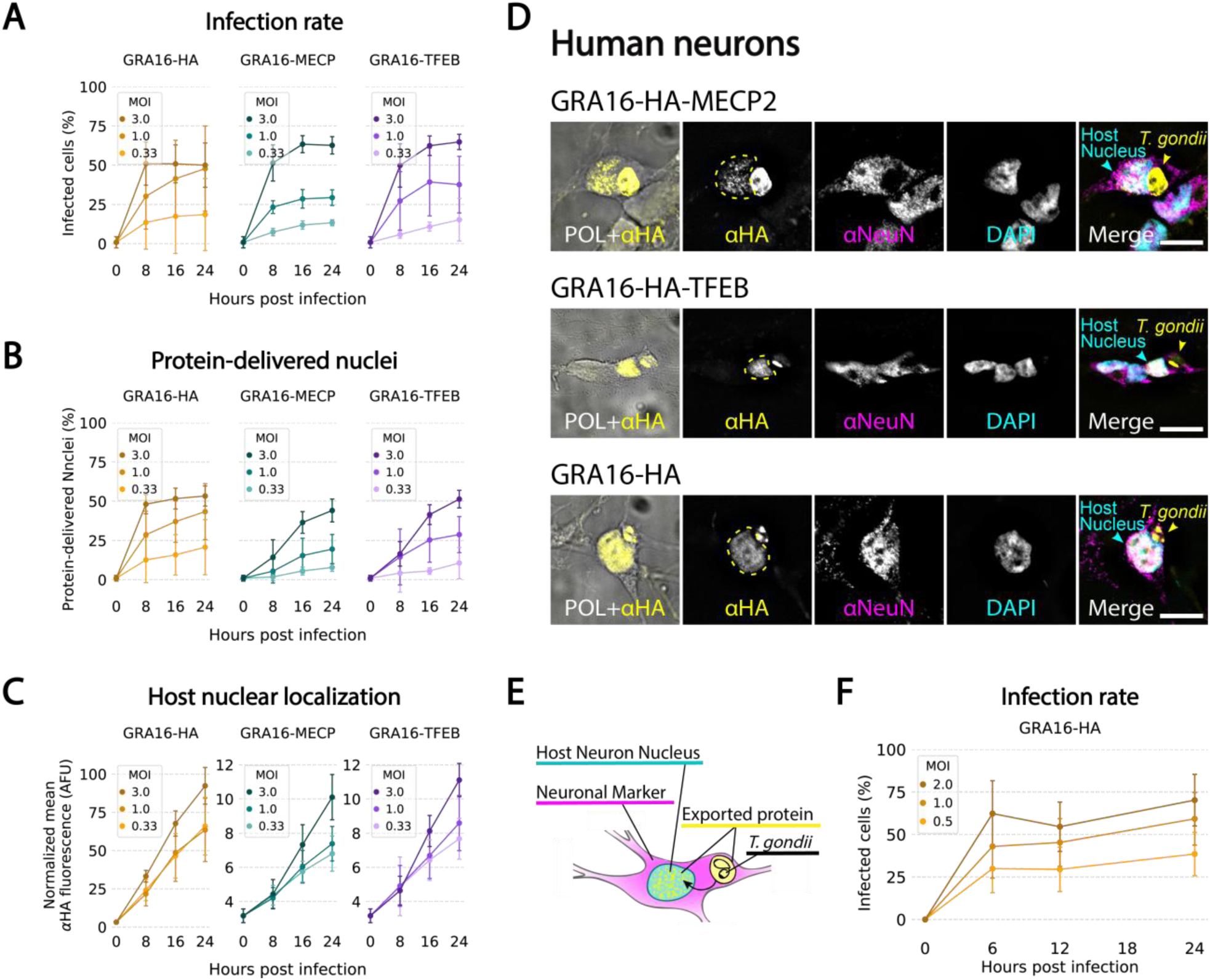
The kinetics of cell infection and protein delivery in human fibroblasts and neurons. (**A**-**C**) Quantitative characterization of the kinetics of cell infection and protein delivery in human foreskin fibroblasts (HFF), infected with different concentrations (MOI) of *T. gondii* expressing GRA16-HA, GRA16-MeCP2 and GRA16-TFEB. (**A**) Infection rate over time. (**B**) Percentage of HFF nuclei labelled positive for the delivered protein over time. (**C**) Normalized mean fluorescence intensity in the nuclei of host cells infected with a single parasite vacuole, over time. Error bars represent mean ±SD. N=24-40 wells per condition (average 1800 cells per well). (**D**) Human neurons differentiated from Lund Human Mesencephalic (LUHMES) cells, infected with *T. gondii* expressing GRA16-HA, GRA16-MeCP2 and GRA16-TFEB. Yellow dashed lines mark the host neuron nucleus. Scale bar: 10µm. (**E**) Illustration of a LUHMES neuron infected with *T. gondii.* Yellow-exported dense-granule protein targeted to the nucleus of the neuron. (**F**) Infection rate in LUHMES neurons infected with *T. gondii* expressing GRA16-HA, over time. Error bars represent mean ±SD. N=10-60 images per condition (average 80 cells per image).

Overall, there was no significant difference in the infection rate of the different lines, indicating that they are similar in their ability to invade host cells (by two-way ANOVA). 24 hours post-administration, the percentage of cells productively infected was on average 50-65% for cells infected using MOI=3, 29-48% with MOI=1 and 13-19% with MOI=0.33. 24 hours post-administration, the mean percentage of host cell nuclei labeled as positive for the proteins was 44-53% for cells infected using a MOI=3, 20-43% with MOI=1 and 8-21% with MOI=0.33. Quantification of fluorescence intensity in the host cell nuclei showed that the fusion of the mammalian proteins MeCP2 and TFEB to GRA16 reduced its accumulation in the host cell nucleus by 7.5 folds on average.

### Delivery of GRA16-fused TFEB and MeCP2 to the nucleus of human neurons

Since the primary targets of protein delivery would be neurons, we tested whether the transgenic *T. gondii* can deliver therapeutic proteins to their site of activity in human neurons. Lund Human Mesencephalic (LUHMES) cells were differentiated *in vitro* to morphologically and biochemically mature dopamine-like neurons (Lotharius et al., 2002; Scholz et al., 2011; Shah et al., 2016) and infected with the *T. gondii* lines GRA16-HA, GRA16-MeCP2 and GRA16-TFEB. All tested lines displayed clear secretion, export and accumulation of the tagged protein in the nuclei of the neurons. Analysis of the kinetics of parasite infection in these neurons showed that 24 hours post-administration, 70% of neurons were infected using MOI=2, 59% using MOI=1 and 39% using MOI=0.5 (Fig. 4D-F).

### *T. gondii*-delivered MeCP2 binds heterochromatic DNA in the nucleus of primary neurons

In order to test the therapeutic relevance of the system, we focused on GRA16-MeCP2. MeCP2 is a nuclear protein which is expressed ubiquitously, but MeCP2 levels are highest in postmitotic neurons, where it is critical for the function and maintenance of the neurons (Lyst & Bird, 2015). This is demonstrated by the disease manifestation of Rett syndrome (RTT), a severe neurological disorder caused by loss-of-function mutations in the *MECP2* gene. The activity of the MeCP2 protein depends on its specific binding of heterochromatic DNA in the nucleus. MeCP2 recognizes patterns of methylated DNA and recruits co-repressor complexes. Therefore, a classical readout for its functionality is its ability to specifically bind heterochromatic DNA (Guy et al., 2018).

To test the functionality of GRA16-fused MeCP2, we infected mouse primary neuronal cultures with *T. gondii* expressing GRA16-MeCP2. In mouse cells, heterochromatic DNA can be readily visualized with DAPI staining. Importantly, we found that the MeCP2 fusion protein delivered by the parasites co-localizes to the foci of heterochromatic DNA in the nucleus (Fig. 5A). This suggests that the MeCP2 fusion protein synthesized and delivered by the parasites binds heterochromatic DNA in the neuronal nucleus, mimicking the functional endogenous MeCP2 (Guy et al., 2018). Similarly, mouse neuroblastoma (N2A) cells infected with *T. gondii* secreting GRA16-MeCP2 presented the same binding of heterochromatic DNA, as well as co-localization with MeCP2 immunostaining. By comparison, cells infected with *T. gondii* parasites secreting GRA16-HA did not present the same co-localization, and the protein was evenly diffused in the nucleus, confirming the specificity of binding to the fusion protein containing MeCP2 (Fig. 5B).

**Fig. 5.**
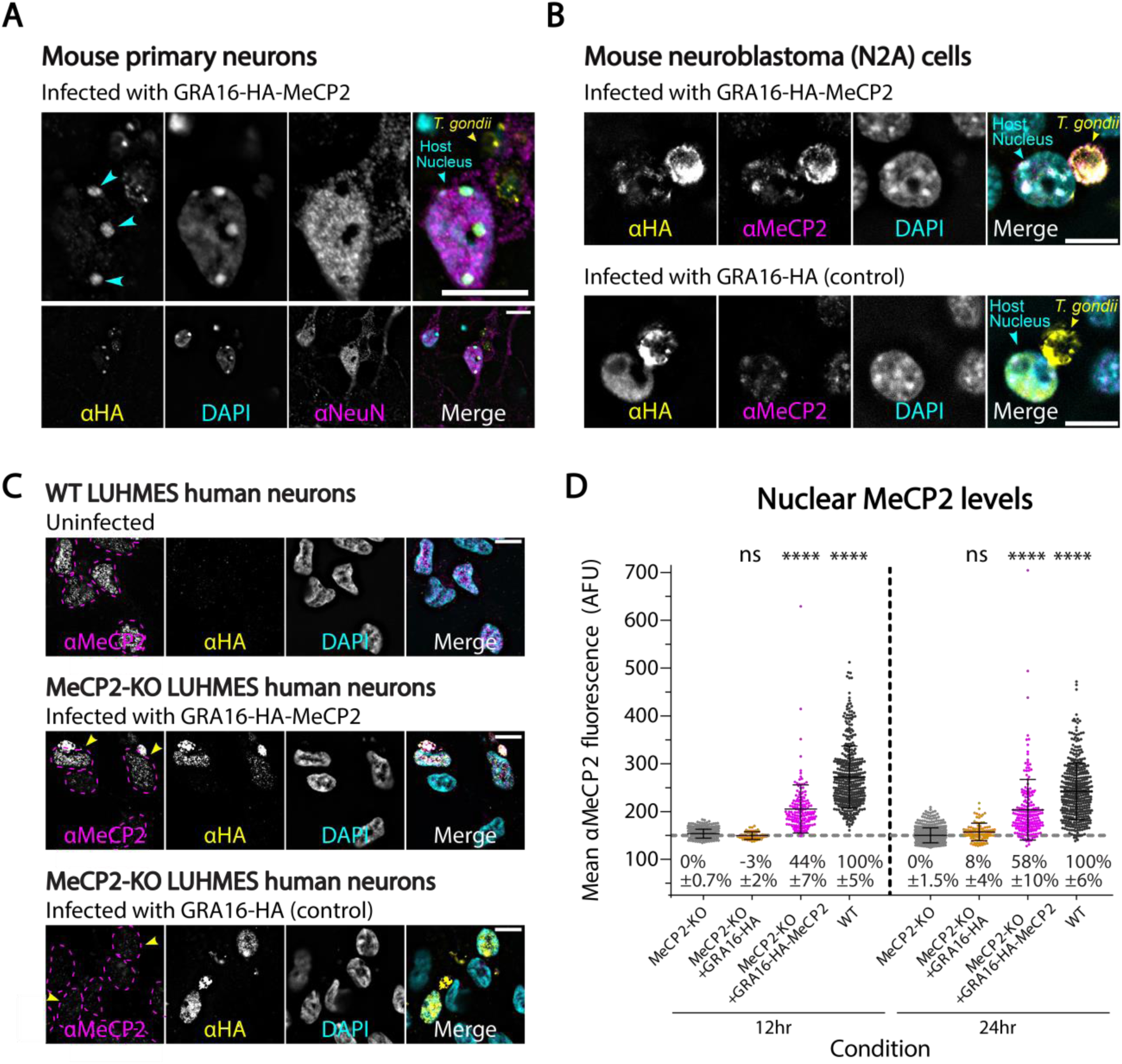
GRA16-MeCP2 binds heterochromatic DNA in neurons and is delivered in therapeutic levels. (**A**) Mouse primary neurons infected with GRA16-MeCP2 *T. gondii*, 12 hours after inoculation. Top images show a close-up of the neuronal soma and intracellular *T. gondii*. Blue arrows mark co-localization of the GRA16-MeCP2 protein with foci of heterochromatic DNA. (**B**) N2A mouse neuroblastoma cells infected with GRA16-MeCP2 and GRA16-HA *T. gondii*, 24 hours after inoculation. (**C**) LUHMES WT and MeCP2-KO human neurons infected with GRA16-MeCP2 and GRA16-HA *T. gondii*, 12 hours after inoculation. Magenta dashed lines mark the neuronal nuclei. Yellow arrows mark infected neurons. (**D**) Fluorescent quantification of the anti-MeCP2 signal in the nucleus of MeCP2-KO neurons, MeCP2-KO neurons infected with GRA16-HA (negative control) or GRA16-MeCP2 *T. gondii*, and WT neurons. Error bars represent mean ±SD. N=50-510 cells per condition. AFU-arbitrary fluorescence units. Text below each scatter plot shows the mean relative nuclear levels of MeCP2 (compared to WT neurons) ±95% confidence interval. Significance represents the difference between each condition and the untreated MeCP2-KO neurons, calculated by two-way ANOVA with multiple comparisons (Dunnett test). **** P<0.0001; ns-not significant. Scale bar: 10µm.

Excess of MeCP2 can be deleterious, as exhibited by the pathogenesis of MECP2 duplication syndrome (Ramocki, Tavyev, & Peters, 2010). Furthermore, gene therapy studies have shown that *MECP2* overexpression leading to 2-6 folds higher protein levels can induce adverse symptoms and increased mortality in mice (Collins et al., 2004; Gadalla et al., 2013; Luikenhuis, Giacometti, Beard, & Jaenisch, 2004). To test the level of MeCP2 rescue achieved by the transgenic *T. gondii*, we quantified MeCP2 delivery in MeCP2-deficient neurons. MeCP2 knock-out (*MECP2*-KO) LUHMES cells (Shah et al., 2016) were differentiated into mature human neurons and infected with GRA16-MeCP2 *T. gondii*. Nuclear levels of MeCP2 in infected and non-infected neurons were quantified by immunofluorescence, and relative protein levels were calculated by comparison to endogenous MeCP2 in wild-type (WT) neurons. We found that 12 hours post-infection *T. gondii*-mediated MeCP2 rescue reaches on average 44% of WT levels, and 24 hours post-infection it reaches 58% of WT levels (P<0.0001, two-way ANOVA). Importantly, MeCP2-KO neurons infected with GRA16-HA *T. gondii* do not show a significant increase in anti-MeCP2 immunostaining (12 hours: P=0.88, 24 hours: P=0.35) (Fig. 5C-D). This suggests that the amount of protein delivered by transgenic *T. gondii* lies within a range compatible with therapeutic benefit (Carrette, Blum, Ma, Kelleher, & Lee, 2018; Gadalla et al., 2013).

## Discussion

In this study we investigated a method for therapeutic intracellular protein delivery via a transgenic brain parasite – *T. gondii*. We assessed different approaches for heterologous protein secretion and characterized *T. gondii* lines that deliver therapeutic proteins to the nuclei of various mammalian cells. We showed that *T. gondii* expressing a GRA16-fused MeCP2 deliver a fusion protein that binds heterochromatic DNA in neurons, mimicking the functional endogenous MeCP2.

For both the Toxofilin and GRA16 fusion proteins, we found that mistargeted proteins were mostly arrested within the secretory pathway (ER, golgi). However, some fusion proteins resulted in unexpected mistargeting to parasite-unique organelles, such as the microneme or apicoplast (**Table S1 and S2**). As the fusion proteins tested in this study represent a wide and unconventional set of sequences, the targeting of the tested fusion proteins could provide an interesting resource to analyze factors affecting protein trafficking in *T. gondii*, as well as the regulation of protein secretion and export from the parasitophorous vacuole (PV).

Interestingly, previous work which attempted to fuse dense granule exported proteins to marker proteins such as fluorescent or other highly structured proteins found that fusion to another protein blocked its ability to be exported through the PV membrane (PVM), causing it to accumulate in the PV space (Curt-Varesano, Braun, Ranquet, Hakimi, & Bougdour, 2016; Hakimi & Bougdour, 2015; Hakimi et al., 2017; Marino et al., 2018). In addition, known exported dense granule proteins of *T. gondii* are enriched in regions of high intrinsic disorder (Hakimi & Bougdour, 2015). In this study we tested the secretion, export and host cell nuclear localization of multiple heterologous proteins fused to the dense granule exported protein GRA16. Of the expressed fusion proteins that were successfully secreted by the dense granules, some were detected in the host cell nucleus (MeCP2, SMN, TFEB) and some were not, suggesting that they were not exported from the PV (ASPA and GALC). When comparing the intrinsic disorder profiles of these proteins, we noticed a significant correlation between the level of protein disorder and the propensity to be exported and localize to the host cell nucleus (Fig. 3, Fig. S1). This provides strong support for the proposed model of protein export, in which high intrinsic disorder is required for protein export through the PVM (Hakimi et al., 2017).

During this study we also established a new model for the study of *T. gondii* in human neurons differentiated from neuronal precursor LUHMES cells. Although for many hosts (including humans), *T. gondii*’s dominant infection is in the CNS, most research on the biology of *T. gondii* is performed in standard fibroblast and epithelial cell lines. On the other hand, use of primary neuronal cultures can be highly variable, expensive, labor intensive and lead to high animal use. LUHMES cells are conditionally-immortalized neuronal precursor cells for which culturing is straightforward and which can differentiate into mature dopamine-like human neurons within one week (Lotharius et al., 2002; Scholz et al., 2011; Shah et al., 2016). This LUHMES neuronal model could facilitate the study of fundamental questions regarding the unique interaction of *T. gondii* with neurons, and especially human neurons.

*T. gondii*-mediated protein delivery has multiple attractive characteristics. *T. gondii* has an exceptionally high CNS-specificity driven by active trafficking to the CNS from peripheral tissues, even with oral administration. Likewise, active motility also drives *T. gondii*’s widespread distribution in the brain – in a systematic characterization of cyst distribution in infected mice, 92% of the brain regions examined were found to contain tissue cysts (Berenreiterová, Flegr, Kuběna, & Němec, 2011). Transgenic *T. gondii* can provide either long-term or transient protein delivery, based on the utilized secretion organelle and subsequent persistence or clearance of the parasites. Importantly, as a biological agent, *T. gondii* is responsive to external cues and can implement engineered genetic circuits (Jimenez-Ruiz, Wong, Pall, & Meissner, 2014; Wang et al., 2016). This could be utilized for incorporating inducible expression systems, or for generating attenuated *T. gondii* which can be cleared from the tissue following protein delivery. *T. gondii* can provide local and compartmentalized protein synthesis *in situ* and deliver proteins directly into the cytosol of cells. It can deliver large proteins, the size limit of which is yet unknown (the largest fusion protein delivered in this study is 110kDa). For comparison, the leading vectors used for gene therapy today, AAV and self-complementary AAV (scAAV), are limited to a packaging capacity of 4.7kb and 2.2-3kb, respectively, which renders them impractical for many human proteins (Colella et al., 2018).

Rett syndrome is a debilitating neurological disorder that arises from mutations in the X-linked *MECP2* gene and affects approximately 1 in 10,000 females. It is characterized by apparently normal early development followed by profound neurologic regression around 1-2 years of age (Carrette et al., 2018; Katz et al., 2016). Since 2007, several gene therapy studies in rodents have shown that expression of functional MeCP2 can reverse the symptoms of Rett syndrome, even after disease onset (Gadalla et al., 2013, 2017; Guy, Gan, Selfridge, Cobb, & Bird, 2007; Katz et al., 2016). Furthermore, exclusive MeCP2 expression in the CNS is sufficient to rescue the majority of Rett syndrome phenotypes (Ross et al., 2016), supporting the therapeutic potential of MeCP2 delivery to the CNS. As a result, several studies are exploring *MECP2* gene therapy, mostly using AAV and scAAV vectors for delivery of the *MECP2* gene (Gadalla et al., 2013; Garg et al., 2013; Katz et al., 2016; Matagne et al., 2017; Sinnett et al., 2017). These studies have been spurring tremendous hope for gene therapy of Rett syndrome.

However, some challenges are still being overcome, such as toxicity from overexpression or mistargeting of the gene, packaging limitations of the viral vectors and limited brain transduction efficiency (Gadalla et al., 2017; Katz et al., 2016). In this study, we demonstrate intracellular delivery of the MeCP2 protein into neurons. Importantly, we also demonstrate that the delivered MeCP2 protein binds heterochromatic DNA in the neurons, mimicking the functional endogenous MeCP2 (Carrette et al., 2018; Gadalla et al., 2013).

In this study we developed and characterized transgenic *T. gondii* lines capable of intracellular delivery of therapeutic proteins based on fusion to GRA16. This approach could be extended to the use of other endogenously-secreted *T. gondii* proteins which exhibit different secretion dynamics and could be adapted to the therapeutic protein of interest. Furthermore, in addition to the delivery of endogenous mammalian proteins, this approach could be used for the delivery of engineered proteins, such as engineered enzymes, ligands, antibodies, transcriptional regulators or programmable nucleases (Gaj, Gersbach, & Barbas, 2013).

## Acknowledgements

We thank all members of the Rechavi and Sheiner labs for their endless help, support, enriching discussions and valuable feedback on the manuscript. We thank Prof. Daniel Frenkel and Prof. Dan Peer from Tel Aviv University for their advice. Dror Cohen from Tel Aviv University for the illustration of figure 1. Dr. Eric Kalkman and Susan Baillie from the Scottish Bioscreening Facility for technical help with the automated high-content imaging systems. Prof. Anita Koshy from the University of Arizona for the Toxofilin-Cre plasmid and Dr. Ruth Shah and Prof.

Adrian Bird from the University of Edinburgh for the LUHMES cell lines. This work was supported by the Nadal-Colton Applied Research Fund, International Rett Syndrome Foundation (Rettsyndrome.org), Adelis Foundation, Glasgow Knowledge Exchange Fund, Naomi Kadar Foundation Fellowship, Joan and Jaime Constantiner Fellowship and Prajs-Drimmer Scholarship. L.S. is a Royal Society of Edinburgh personal research fellow.

## Author Contribution

S.B., O.R. and L.S. conceived the study and designed the experiments, S.B. and K.H. performed the experiments and analyzed the data, S.B. prepared the figures and performed the formal analysis, P.D.R. and S.C. helped with the MeCP2 experiments and provided the primary neurons, S.B. wrote the manuscript with input from O.R., L.S., S.C., K.H. and P.D.R.

## Data and material accessibility

Raw microscopy files, metadata, code and data tables are available on: https://github.com/shaharbr.

## Supplementary Materials

**Fig. S1.**
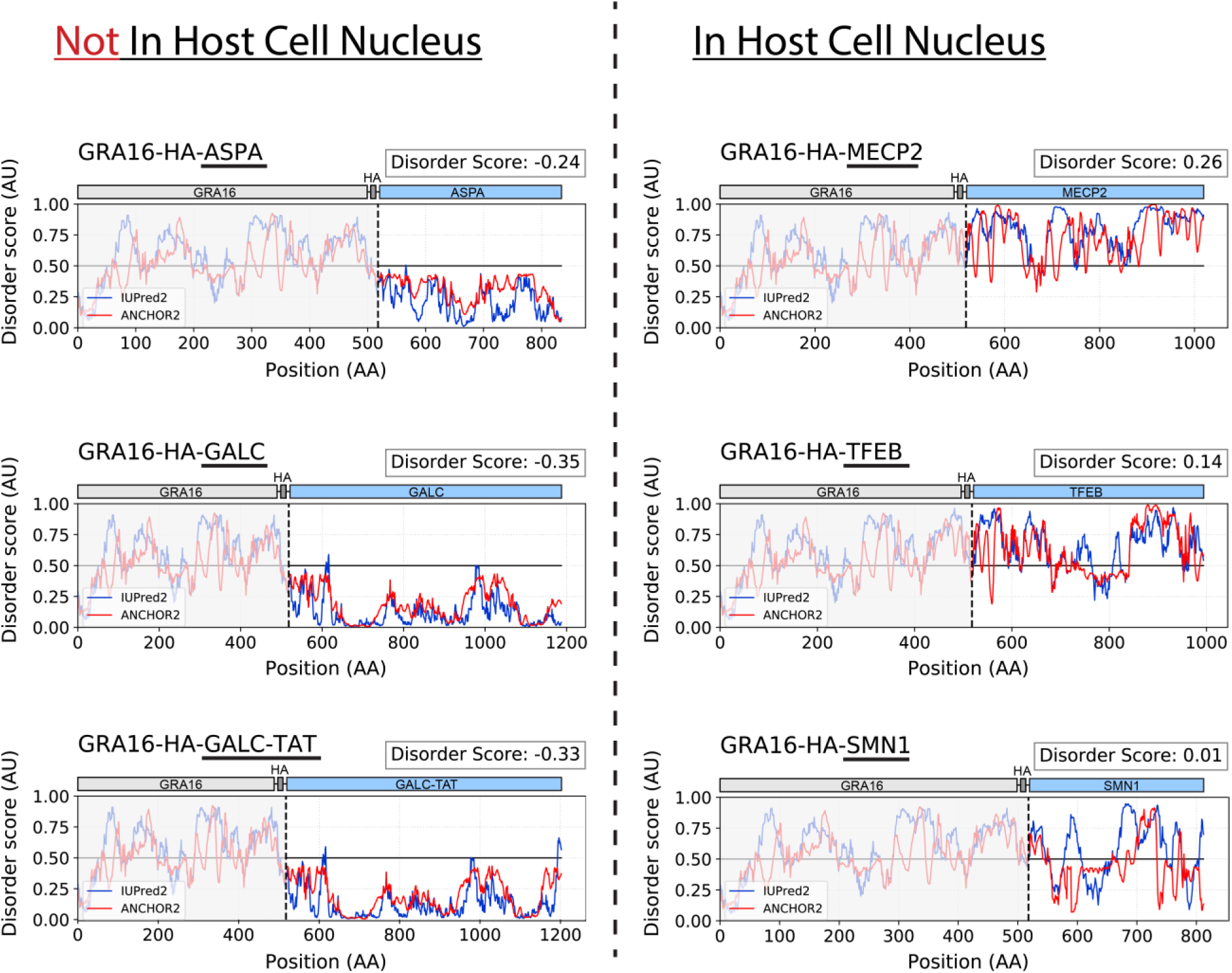
The level of intrinsic disorder of the GRA16-fused therapeutic proteins predicts their localization to the host nucleus. Prediction of protein intrinsic disorder of GRA16 fusion proteins, using IUPred2 (disordered protein regions) and ANCHOR2 (disordered binding regions). Grey: GRA16-HA. Dashed lined marks the start of the heterologous protein. Disorder score = average(IUPred2 score, ANCHOR2 score) of the fused therapeutic protein.

**Fig. S2.**
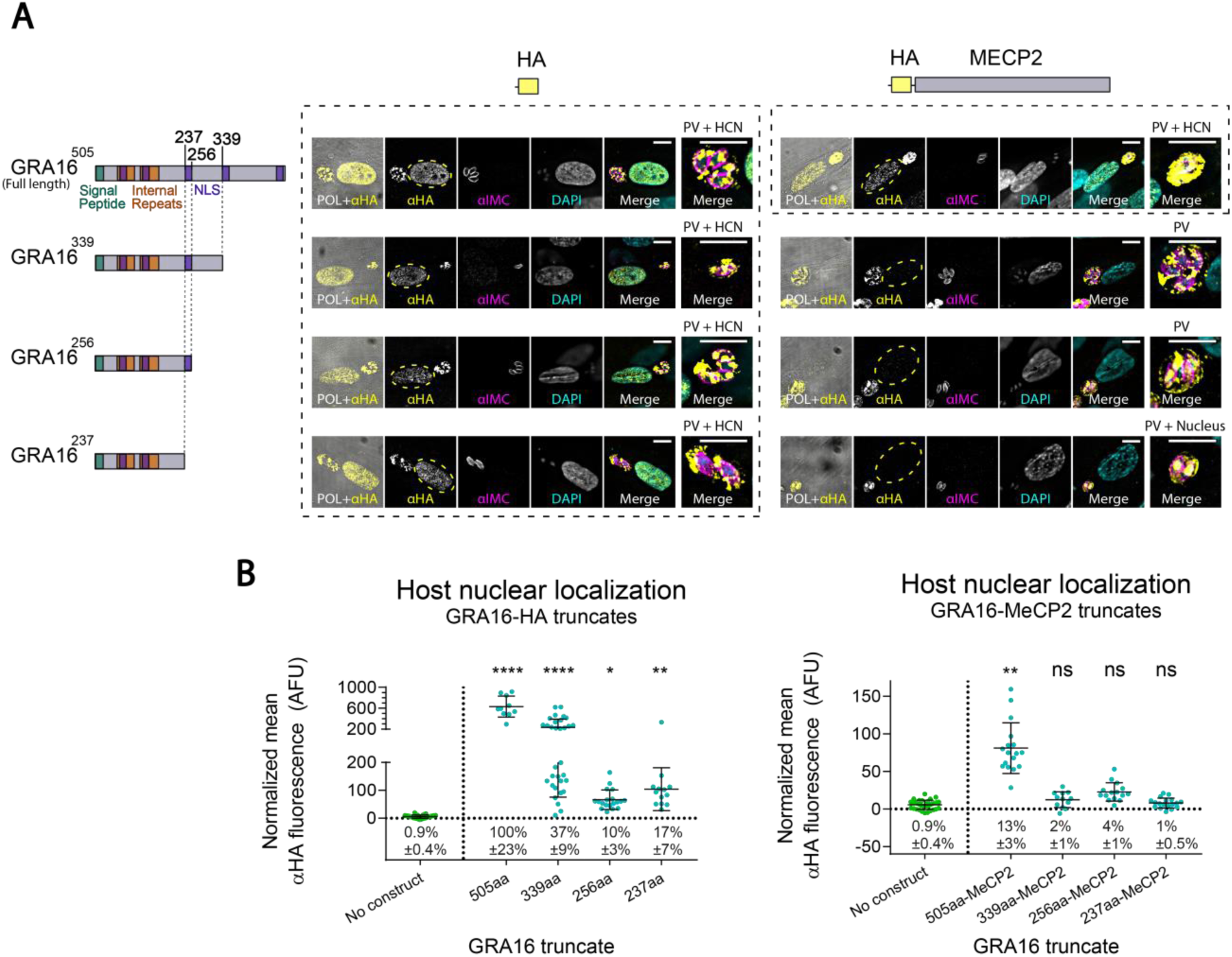
Truncates of GRA16 do not deliver MeCP2 to the host nucleus. (**A**) Intracellular *T. gondii* stably expressing different fragments of GRA16, fused to either a HA tag alone or to an HA-tagged murine MeCP2 (codon optimized). All tagged proteins localized to the PV. However, only the GRA16 truncates without MeCP2 localized to the host cell nucleus (HCN), as well as the full-length GRA16 fused to MeCP2. Rightmost image of each set shows a close-up of the parasite vacuole, and the text above it indicates the localization of the tagged protein in the image. Yellow dashed lines mark the host cell nucleus. Top and left: illustration of the full-length GRA16 (Bougdour et al., 2013)), C-terminally truncated GRA16, and the fused HA or HA-MECP2 sequences used in the respective constructs. NLS-Nuclear localization signal. (**B**) Fluorescence quantification of the anti-HA signal in the nucleus of infected host cells. Error bars represent mean ±SD. N=10-32 cells per condition. Take note that the y-axis of the leftmost graph is divided to two segments, to account for the high values recorded from GRA16-HA. AFU-arbitrary fluorescence units. Text below each scatter plot shows the mean relative nuclear localization ±95% confidence interval (relative to the full-length GRA16[505]-HA). Significance represent the difference between the protein and the parental strain (‘no construct’), calculated by two-way ANOVA with multiple comparisons (Dunnett test). **** P<0.0001; ** P<0.0021; * P<0.0332; ns-not significant. Scale bar: 10µm.

**Table S1.**
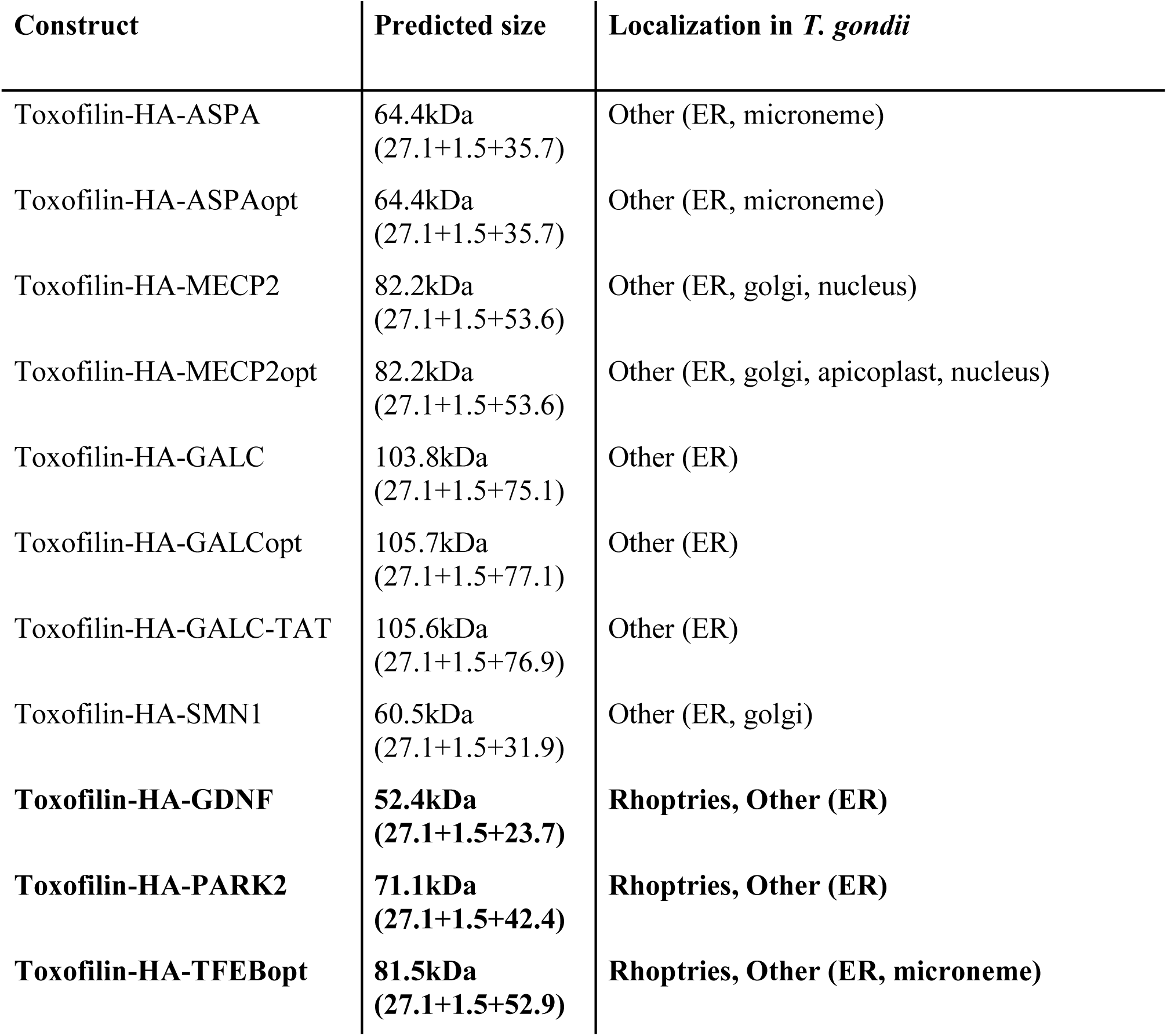
Toxofilin fusion constructs used in this study. Right column summarized the intracellular localizations in *T. gondii* observed for each construct. Constructs that presented correct localizations to the targeted rhoptry secretion organelle are highlighted in bold text.

| Construct | Predicted size | Localization in *T. gondii* |
| --- | --- | --- |
| Toxofilin-HA-ASPA | 64.4kDa<br>(27.1+1.5+35.7) | Other (ER, microneme) |
| Toxofilin-HA-ASPAopt | 64.4kDa<br>(27.1+1.5+35.7) | Other (ER, microneme) |
| Toxofilin-HA-MECP2 | 82.2kDa<br>(27.1+1.5+53.6) | Other (ER, golgi, nucleus) |
| Toxofilin-HA-MECP2opt | 82.2kDa<br>(27.1+1.5+53.6) | Other (ER, golgi, apicoplast, nucleus) |
| Toxofilin-HA-GALC | 103.8kDa<br>(27.1+1.5+75.1) | Other (ER) |
| Toxofilin-HA-GALCopt | 105.7kDa<br>(27.1+1.5+77.1) | Other (ER) |
| Toxofilin-HA-GALC-TAT | 105.6kDa<br>(27.1+1.5+76.9) | Other (ER) |
| Toxofilin-HA-SMN1 | 60.5kDa<br>(27.1+1.5+31.9) | Other (ER, golgi) |
| **Toxofilin-HA-GDNF** | **52.4kDa**<br>**(27.1+1.5+23.7)** | **Rhoptries, Other (ER)** |
| **Toxofilin-HA-PARK2** | **71.1kDa**<br>**(27.1+1.5+42.4)** | **Rhoptries, Other (ER)** |
| **Toxofilin-HA-TFEBopt** | **81.5kDa**<br>**(27.1+1.5+52.9)** | **Rhoptries, Other (ER, microneme)** |

**Table S2.** GRA16 fusion constructs used in this study. Right column summarized the intracellular localizations in *T. gondii* observed for each construct. Constructs that presented localizations to the PV and host cell nucleus are highlighted in bold text.

| Construct | Predicted size | Localization in *T. gondii* |
| --- | --- | --- |
| GRA16-HA-ASPA | 92.2kDa<br>(54.9+1.5+35.7) | PV, Other (ER, golgi, nucleus) |
| GRA16-HA-ASPAopt | 92.2kDa<br>(54.9+1.5+35.7) | PV, Other (ER, golgi, nucleus) |
| GRA16-HA-GALC | 131.6kDa<br>(54.9+1.5+75.1) | PV, Other (ER) |
| GRA16-HA-GALCopt | 133.5kDa<br>(54.9+1.5+77.1) | Other (ER) |
| GRA16-HA-GALC-TAT | 133.3kDa<br>(54.9+1.5+76.9) | Not expressed |
| **GRA16-HA-MECP2opt** | **110kDa**<br>**(54.9+1.5+53.6)** | **PV + Host Cell Nucleus, Other (ER, golgi)** |
| **GRA16-HA-SMN1** | **88.3kDa**<br>**(54.9+1.5+31.9)** | **PV + Host Cell Nucleus, Other (ER, golgi, apicoplast, microneme)** |
| **GRA16-HA-TFEBopt** | **109.3kDa**<br>**(54.9+1.5+52.9)** | **PV + Host Cell Nucleus, Other (ER)** |

## Materials and methods

### *T. gondii* culture and maintenance

Type I RH *T. gondii* were grown in human foreskin fibroblasts (HFF) in high glucose Dulbecco’s modified Eagle’s medium (DMEM) supplemented with 4 mM L-glutamine, 10% Fetal Bovine Serum (FBS) and 1% penicillin/streptomycin or 20 µg/ml gentamicin antibiotics, thereby referred to as “complete DMEM”. Cultures were monitored daily, and the parasites were passaged by transferring a drop (20-100 µl) of the supernatant of a lysed dish (containing extracellular parasites) into a fresh dish with confluent HFF cells.

### DNA transfection of *T. gondii*

Tachyzoites of *T. gondii* were collected from the supernatant of lysed cells or mechanically released by scraping and passing through a 23-26 gauge needle. Extracellular parasites were filtered using a 3µm-pore filter and pelleted by centrifugation at 800G for 5 minutes. Supernatant was discarded, and parasites were resuspended in cytomix buffer (10 mM K2HPO4/KH2PO4, 25 mM HEPES, 2 mM EGTA pH 7.6, 120 mM KCl, 0.15 mM CaCl2, 5 mM MgCl2 with 5 mM KOH adjusted to pH 7.6) freshly supplemented with 3 mM ATP and 3 mM GSH. Resuspended parasites in a total volume of 350 µl (for the BTX system) or 800 µl (for the Bio-Rad system) were placed in an electroporation cuvette and supplemented with up to 50 µl of the transfected DNA (10-80 µg DNA). The DNA constructs were transfected into the parasites as either circular plasmids or after linearization with ScaI. The parasites were electroporated using two square-wave pulses. On the BTX ECM 830 square wave electroporator, we used the High Voltage (HV) mode and the following settings: voltage 1700 V, pulse length 0.05 milliseconds, 2 pulses, 200 milliseconds interval, unipolar. On the Bio-Rad GenePulser Xcell electroporator, we used the square wave protocol (program 3) with the following settings: voltage 1700 V, pulse length 0.2 milliseconds, 2 pulses, 5 seconds interval, 4 mm cuvette. The transfected parasites were transferred onto cells seeded on glass coverslips. The cells were fixed 24 and 48 hours after infection, immunofluorescently stained and microscopically assessed for transfection efficiency, transient protein expression (expression from extrachromosomal DNA) and localization.

### Generation of *T. gondii* clonal lines

To generate stable clonal lines, *T. gondii* were transfected as described, and genomic integration of the exogenous DNA was selected for by drug selection followed by clone isolation. After transfection, parasites were transferred onto a fresh dish with HFF cells. The next day, media was changed to fresh media containing the drug used for selection (pyrimethamine for DHFR-TS selection or mycophenolic acid+Xanthine for HXGPRT positive selection).

Starting from the first detection of egressed extracellular parasites in the culture supernatant, 50-200 µl drops of the supernatant were passed daily to a second dish of HFF cells in selective media. When the parasites in the second dish started egressing from the HFF, drops of the supernatant were passed daily to a third dish, and so on. After the 3rd-4th dish started lysing (about 3 weeks after transfection), the parasites were considered a “stable pool” containing parasites that integrated the exogenous DNA construct into their genome. The stable pool was immunofluorescently stained to assess the percentage of construct-expressing parasites in the stable pool as well as to evaluate protein localization upon genomic integration of the construct. Clonal lines were isolated in 96-well plates with HFF by limiting dilutions or by FACS sorting single parasites into each well. After 5-10 days of growth, clones were selected as wells containing a single plaque, and tested for integration of the construct via PCR using genomic lysate as template. Genomic lysates were prepared in lysis buffer (10% proteinase K in TE buffer) by incubation at 60 °C for 60 minutes followed by 95 °C for 10 minutes. PCR-positive clones were verified by immunofluorescent staining. Protein localization was determined morphologically based on comparison to the polarized light image, and to co-staining with DAPI and anti-IMC1. Each fusion protein localization was validated over 2-5 independent transfections. At least 100 parasites were microscopically assessed by eye per transfection, and at least 20 z-stack images were acquired for each transfection, for transient and for stable expression (**Tables S1 and S2**).

### Culture, neuronal differentiation and infection of LUHMES cells

Culturing and differentiation of LUHMES cells were carried according to Shah et al. (Shah et al., 2016). Undifferentiated and differentiated LUHMES cells were grown in 75cm2 flasks coated overnight with fibronectin+Poly-L-ornithine (1 µg/µl fibronectin and 43µg/ml PLO in sterile distilled water). Undifferentiated LUHMES cells were split every 3 days and seeded at 106 cells per 75cm2 flask. They were maintained in a proliferation media consisting of advanced DMEM/F12 media supplemented with 1x N2 serum-free supplement, 2 mM L-glutamine and 40 ng/ml beta-FGF. LUHMES cell were differentiated into morphologically and biochemically mature dopamine-like neurons using a differentiation media consisting of advanced DMEM/F12 media supplemented with 2 mM L-glutamine, 1x N2 serum-free supplement, 1 µg/ml tetracycline, 2 ng/ml GDNF and 1 mM cell-permeable cAMP analog (N6,2′-O-Dibutyryladenosine 3′,5′-cyclic monophosphate sodium salt). A day before initiating differentiation (day -1), the cells were seeded at a higher density of 2.5*106 cells per flask in normal proliferation media. The next day (day 0), differentiation was initiated by changing the media to the differentiation media. 2 days later (day 2), the cells were dissociated using trypsin and seeded in new flasks at a density of 6*106 cells per flask, or on coated glass coverslips in 24-well plates at 0.3*106 cells per well, in fresh differentiation media. On day 6 of differentiation (at which point the cells are mature neurons, as confirmed by anti-NeuN staining), tachyzoites of the selected *T. gondii* line were resuspended in differentiation media and administered to the neurons.

### Culture and infection of N2A cells

N2A (Neuro2A, mouse neuroblastoma) cells were cultured in high glucose DMEM supplemented with 10% FBS, 1% penicillin/streptomycin, 2 mM L-glutamine and 0.1 mM MEM non-essential amino acids solution. The day before infection, they were seeded on glass coverslips. The next day, tachyzoites of the selected *T. gondii* lines resuspended in the N2A media and administered to the N2A cells at MOI=1. The cells were fixed 24 hours after infection, and immunofluorescently stained as described.

### Preparation and infection of mice primary neuronal cultures

The cortex and hippocampi of WT P1 pups were dissociated enzymatically using papain and mechanically using gentle pipettation. Cells were counted and seeded in a 24-well plate on poly-l-lysin coated glass coverslips at a density of 100,000 cells per well, in neurobasal media (NBA, 2% B27 and 1% L-glutamine). After 5 days in culture, cells were infected with tachyzoites of the GRA16-MeCP2 *T. gondii* line at MOI=1. 12 hours post inoculation, cells were washed with PBS, fixed and immunofluorescently stained as described.

### Immunofluorescent staining (Immunofluorescence assay, IFA)

Cells were grown on 13 mm glass coverslips in a 24-well plates. At the respective time-point, cells were fixed in 4% PFA for 20 min at room temperature (RT). They were permeabilized and blocked by incubation with 2% blocking solution (2% bovine serum albumin, 0.2% Triton-X in PBS) for 20 minutes at RT. Blocking solution was removed, coverslips were covered with primary antibodies in blocking buffer and incubated for 1 hour at RT. Following 3 washes with PBS (5 minutes, shaking), they were covered with secondary antibodies in blocking buffer and incubated for 40 minutes at RT, protected from light. Coverslips were washed in PBS 3 times, dipped briefly in water, blotted on paper to remove excess water and mounted on slides with Fluoromount-G mounting media containing DAPI (Southern Biotech, 0100-20). Slides were allowed to dry overnight at RT prior to imaging. For long term storage, slides were kept at 4 °C in the dark.

### Microscopy data acquisition and analysis

Unless specified otherwise, all slides were imaged using a DeltaVision Core microscope (AppliedPrecision) using a 100x objective. Images were handled using the Fiji distribution of ImageJ, imported using the OME bio-formats plugin (Linkert et al., 2010) and deconvolved using the Diffraction PSF 3D and Iterative Deconvolution 3D plugins (Dougherty, 2005). All adjustments of brightness and contrast were linear and applied to the entire image equally.

Background fluorescence was determined by sampling an “empty” area of the image, and maximum display threshold for the image was set to allow optimal visualization of the cell structures and protein localization (e.g. in the host cell nucleus). All raw images, imaging metadata, and ImageJ macros detailing the processing protocol are available on: https://github.com/shaharbr.

### Automated high-content imaging of the kinetics of *T. gondii* infection and protein secretion in HFF cells

HFF cells were seeded in five 384-microwell plates corresponding to five fixation time points (8, 16, 24, 32, 40 hours), at a dilution of 3,500 cells per well in 50 µl complete DMEM and given 2 days to reach confluency (calculated 4,500 cells per well). Tachyzoites of the lines GRA16-HA, GRA16-MeCP2 and GRA16-TFEB were syringe-egressed, filtered and counted in a Neubauer chamber haemocytometer, and diluted to the appropriate concentration. Wells were infected with MOI (multiplicity of infection) of 3, 1 or 0.33 (13,500, 4,500 or 1,500 parasites in 20 µl per well, respectively), or received media only (“MOI 0”). At each timepoint, the respective plate was washed with PBS and fixed manually with 4% PFA. Each condition (parasite line+MOI+timepoint) was repeated over 32-40 wells. Immunostaining was performed using a Beckman Coulter Biomek FXp liquid handling robot with a Thermo Multidrop Reagent Dispenser and with a MANTIS Liquid Handler. Cells were first permeabilized and blocked with 2% blocking solution for 20 min at RT. Blocking buffer was removed and cells were incubated with 5 µl per well of the primary antibody solution (anti-HA + anti-IMC1) in blocking solution, for 60 min at RT. Cells were washed 3 times with PBS, and then incubated with 20 µl secondary antibody solution and Hoechst 33342 dye (diluted 1:50,000) in blocking solution, for 40 min at RT and protected from light. Cells were washed 3 times with PBS. Plates were then imaged on the GE IN Cell 2000 platform. 5 fields-of-view were acquired from each well (average 360 cells per field-of-view).

### Manual imaging of the kinetics of *T. gondii* infection and protein secretion in LUHMES neurons

LUHMES cells were differentiated to neurons in 24-well plates with coated glass coverslips using the protocol described above. On day 6 of differentiation, tachyzoites of the line GRA16-HA were administered at MOI 0.5, 1 or 2. On each timepoint (6, 12 24 and 32 hours post-infection), the respective wells were washed with PBS, fixed and immunofluorescently stained with anti-HA and anti-IMC1 or anti-HA and anti-NeuN manually as described. Each condition (MOI+timepoint) was repeated over 3 coverslips. 10-60 random regions-of-interest (average 80 neurons per image) were imaged for each coverslip on the BX63 Olympus microscope using a 40x objective.

### Analysis of *T. gondii* infection and protein delivery kinetics in HFF and in LUHMES neurons

Image analysis was performed using the open-source CellProfiler (Carpenter et al., 2006) software. Full protocol used for the image analysis, including the parameters chosen for identification of host cells and intracellular *T. gondii*, is available together with the raw data and image analysis outputs on: https://github.com/shaharbr. In brief, host cell nuclei were identified using the DAPI channel and *T. gondii* were identified using the 594nm channel (corresponding to anti-IMC1 staining). Each *T. gondii* vacuole was associated to a host cell nucleus based on proximity. Fluorescence intensity on the 488nm channel, corresponding to the anti-HA staining was used to quantify the levels of GRA16 fusion-protein localization in the parasite vacuoles and in the host cell nuclei. Importantly, timepoints above 24 hours post infection were removed from the dataset as we found that the parasite vacuoles were too large for efficient segmentation and host cell association by the used image analysis tool. Resulting data tables were analyzed using a custom python code. In brief, this analysis involved labeling and organization of the data, removal of images in which parasite identification failed, removal of outlier wells with exceptionally high fluorescence intensity (>5 std from mean of condition), normalization by subtraction of background fluorescence, labeling of cells infected with a single parasite vacuole, labeling of nuclei positive for the tagged protein (threshold set as above 99% of uninfected cells), calculating descriptive statistics for each well (N= number of cells) and for each condition (N= number of wells) and plotting. Full code and analysis outputs are available on: https://github.com/shaharbr.

### Measuring host nuclear fluorescence intensity in HFF and neurons

Cells infected with *T. gondii* expressing the respective construct were fixed at the designated time points (24 hours for HFF, 12-24 hours for LUHMES), and immunofluorescently stained as described. Infected HFF were stained with anti-HA and anti-IMC1, and infected LUHMES neurons were stained with anti-HA and anti-MeCP2 (for quantifying nuclear MeCP2 in infected neurons) or with anti-NeuN and anti-MeCP2 (for verifying differentiation), and imaged. Mean nuclear intensity of the respective fluorescent signal was measured using imageJ’s ‘Measure’ function.

For HFF, each infected cell nucleus was measured alongside 3 random background regions in the same image. To calculate the normalized mean nuclear intensity, the average of the three background regions was subtracted from the value recorded from the infected cell nucleus, and divided by the exposure time (to account for different exposure times used in imaging of the different transfections). Relative nuclear localization of the expressed protein was calculated by dividing each normalized intensity measurement by the mean normalized intensity of GRA16-HA.

For LUHMES neurons, all nuclei detected in each image were measured. For each measured nucleus, it was labeled whether the neuron contained a *T. gondii* vacuole, and whether the nucleus was also HA-positive. Neurons that contained a *T. gondii* vacuole and were HA-positive were considered ‘infected’ and neurons that did not contain a *T. gondii* vacuole and did not present nuclear anti-HA staining were considered ‘uninfected’. Relative nuclear localization of MeCP2 was calculated by dividing each normalized intensity measurement by the mean normalized intensity of WT neurons (endogenously expressing hMeCP2) from the same timepoint.

### Intrinsic disorder score and correlation to nuclear localization of the fusion proteins

The intrinsic disorder profile for the full translated open read frames of each GRA16 fusion protein was calculated using https://iupred2a.elte.hu/, using the following parameters: Prediction type: IUPred2 long disorder (default), Context-dependent prediction: ANCHOR2 (Mészáros et al., 2018). A custom python code was used to calculate the averaged intrinsic disorder score of the fused therapeutic protein, calculate the correlation between the intrinsic disorder score and nuclear localization of each fusion protein and generate plots. The averaged intrinsic disorder score was calculated as the average of the ANCHOR2 score and the IUPred2 score along the span of the fused therapeutic protein (excluding GRA16-HA), minus the 0.5 threshold for disorder. Protein disorder data and code are available on: https://github.com/shaharbr.

### Molecular cloning

For the plasmids encoding for Toxofilin-fused proteins, we used as a backbone a pGRA vector containing *Toxofilin* cDNA fused to Cre recombinase, surrounded by the 1.1 kb genomic sequence upstream to *Toxofilin* (‘Toxofilin promoter’) and the 3’ UTR of *GRA2*, a kind gift from

A. A. Koshy (Koshy et al., 2010; Lodoen et al., 2010). To generate the therapeutic fusion constructs, we used the following mammalian cDNA sequences: human *SMN1* (Addgene, #37057), human *ASPA* (DNASU, HsCD00044152), *ASPAopt* (GenScript, custom synthesis, human *ASPA* codon optimized for *T. gondii*), human *PARK2* (MGC, BC022014), human *GDNF* (MGC, BC069119), murine *MECP2* isoform 1e/B (GenScript, OMu23690), *MECP2opt* (GenScript, custom synthesis, murine MeCP2 isoform 1e/B codon optimized for *T. gondii*), *TFEBopt* (GenScript, custom synthesis, human TFEB isoform 1 codon optimized for *T. gondii*), human *GALC* (MGC, BC036518), *GALCopt* (GenScript, custom synthesis, human *GALC* isoform 1 [NP_000144] codon optimized for *T. gondii*).

To generate plasmids for the expression of *Toxofilin*-fused *GDNF*, *PARK2* and *MECP2*, we PCR-amplified the mammalian cDNA with primers that add an EcoRV restriction site at the 5‘ and a PacI restriction site at the 3’ of the cDNA. The Cre recombinase was replaced with the amplified mammalian cDNA by restriction-ligation (using SfoI and PacI for the backbone and EcoRV and PacI for the cDNA, SfoI and EcoRV both produce blunt ends).

To generate plasmids for the expression of *Toxofilin*-fused *ASPA*, *SMN1*, *GALC*, *GALC-TAT* and *MECP2opt*, we amplified the pGRA vector and cDNA, and used NEBuilder assembly. As *GALC* contains a region of exceptionally high %GC at the start of the gene, PCR amplification and NEBuilder assembly resulted in a deletion at the start of the gene. To fix this, we performed an additional cloning step to add the missing fragment using the ‘round-the-horn’ (Stephen Floor, 2018) PCR blunt cloning method for both the *Toxofilin-GALC* and *Toxofilin-GALC-TAT* plasmids.

To generate plasmids for the expression of Toxofilin-fused *TFEBopt*, *ASPAopt* and *GALCopt*, we replaced the *MEP2opt* in the Toxofilin-MECP2opt plasmid with the respective cDNA using restriction-ligation with EcoRV and PacI which were added to the sequence during DNA synthesis.

To generate plasmids for the expression of *GRA16*-fused genes, *GRA16* and its promoter were amplified from RH *T. gondii* genomic DNA (primers were designed based on (Bougdour et al., 2013)). These *GRA16* PCR products were used to replace the Toxofilin promoter and coding sequence in the *Toxofilin-ASPA* plasmid, with the *GRA16* promoter and coding sequence, to produce the *GRA16-ASPA* plasmid (by NEBuilder assembly).

To generate plasmids for the expression of *GRA16*-fused *MECP2opt*, *SMN1*, *GALC*, *GALC-TAT*, *TFEBopt*, *GALCopt* and *ASPAopt*, we replaced the *ASPA* with the respective cDNA using restriction-ligation with EcoRV+PacI.

*GRA16* truncate vectors (*HA*-fused and *HA-MECP2*-fused) were generate from the GRA16-HA and GRA16-MECP2opt vectors using the ‘round-the-horn’ method.

### Antibodies

Antibodies and the respective concentrations they were used in, are as follows: anti-HA (Sigma-Aldrich ROAHAHA/Roche clone 3F10, 1:1000), anti-IMC1 (gift from Prof. Dominique Soldati-Favre, 1:2000), anti-MeCP2 (Cell Signaling #3456, 1:200), anti-NeuN (Abcam ab104224, 1:500), Alexa Fluor Goat anti-Rat 488 and 594 (Invitrogen #A-11006 and #A-11007, 1:1000), Alexa Fluor Goat anti-Rabbit 488 and 594 (Invitrogen #A-11008, #A-11012, 1:1000), Alexa Fluor anti-mouse 488 and 594 (Invitrogen #A-11001 and #A-11005, 1:1000).

### Statistical analysis

To determine the significance of nuclear localization of each GRA16-fused protein, we performed one-way ANOVA for the effect of the expressed construct, with multiple comparisons to the “no construct” control. To determine the significance of nuclear localization of each truncated variant of GRA16, fused to HA or to HA-MeCP2, we performed two-way ANOVA for the effect of the GRA16 variant (rows) and of the fused sequence (HA or HA-MECP2, columns), with multiple comparisons to the “no construct” control. To determine the significance of the difference in infection rate between parasite lines in the kinetics analysis, for each timepoint we performed two-way ANOVA for the effect of parasite line (columns) and MOI (rows). To determine the significance of nuclear levels of MeCP2 and GRA16-MeCP2, we performed two-way ANOVA for the effect of the time after inoculation (rows) and the condition (columns), with multiple comparisons to the to the “uninfected *MECP2*-KO” control. Multiple comparisons were performed using the following parameters: within each row, compare the means of each group to control, report multiplicity adjusted P value for each comparison (Dunnett test), one family per row.

